# Targeted Acidosis Mediated Delivery of Antigenic MHC-Binding Peptides

**DOI:** 10.1101/2023.10.18.562409

**Authors:** Joey J. Kelly, Emily Ankrom, Damien Thévenin, Marcos M. Pires

## Abstract

Cytotoxic T lymphocytes are the primary effector immune cells responsible for protection against cancer, as they target peptide neoantigens presented through the major histocompatibility complex (MHC) on cancer cells, leading to cell death. Targeting peptide-MHC (pMHC) complexes offers a promising strategy for immunotherapy due to its specificity and effectiveness against cancer. In this work, we exploit the acidic tumor micro-environment to selectively deliver antigenic peptides to cancer cells using pH(low) insertion peptides (pHLIP). We demonstrated that the delivery of MHC binding peptides directly to the cytoplasm of melanoma cells resulted in the presentation of antigenic peptides on MHC, and subsequent activation of T cells. This work highlights the potential of pHLIP as a vehicle for targeted delivery of antigenic peptides and their presentation *via* MHC-bound complexes on cancer cell surfaces for activation of T cells with implications for enhancing anti-cancer immunotherapy.

## Introduction

Cytotoxic effects associated with conventional cancer treatments have significantly limited their overall effectiveness.^1^ In addition, off-target effects also pose challenges in terms of dosing and administration by causing unwanted side effects to normal cells.^2^ Balancing the therapeutic benefits with the risk of toxicity is critical in minimizing damage to healthy tissue and to improving existing treatments. To overcome these limitations, there is an increasing demand for innovative cancer treatment strategies that can overcome the cytotoxicity-associated drawbacks and improve pharmacological properties.

An effective method to mitigate off-target effects involves linking therapeutic agents to well-defined carriers that precisely target cancer cells. Current advances in targeted cancer treatment primarily leverage the overexpression of specific biomarkers on cancer cell surfaces to administer high doses of therapeutic compounds more precisely.^3^ One prominent example of this class of pharmaceuticals is antibody-drug conjugates (ADCs), which increase the local concentration of chemotherapeutic drugs in cancerous cells while reducing the toxicity to nearby healthy tissue. An important consideration in designing these types of therapeutics is the specific covalent attachment used to link the therapeutic payload.^4^ Numerous covalent linkers, including peptides, disulfides, and thioethers, have been employed to conjugate therapeutic payloads to the antibody.^5–9^ Additionally, other delivery systems such as nanoparticles, liposomes, and dendrimers have been engineered to target overexpressed biomarkers such as epidermal growth factor receptors, folate receptors, surface glycoproteins, and transferrin receptors to effectively target cancer.^3, 10–14^ While targeting overexpressed biomarkers has been successful in increasing the effective concentration of therapeutic compounds in cells, the same biomarkers expressed at low levels on healthy cells have contributed to significant levels of off-target toxicity, highlighting the need to develop alternative strategies.^15^

Another strategy to enhance targeted delivery toward tumors involves exploiting the acidic microenvironment that is characteristic of solid tumors. Most tumors display rapid growth levels, which demands increased energy production via glycolysis. This metabolic shift produces lactic acid as a byproduct and the expulsion of protons from the cancer cells into the extracellular space, a phenomenon known as the Warburg effect.^16^ As a result, the extracellular space surrounding cancer cells typically has an acidic pH ranging from 6.7 to 7.1 as opposed to a healthy pH range of 7.35 to 7.45. Interestingly, the environment closest to the cell surface is even more acidic; with the pH reaching 6.1, making them an ideal target for pH(low) insertion peptides (pHLIPs).^17^ The distinctive feature of pHLIP centers on its ability to specifically target the acidic microenvironment by undergoing a pH-dependent rearrangement. This pH-dependent change leads to the insertion of its C-terminus across the cell membrane, forming a transmembrane α-helix.^18, 19^ Importantly, pHLIP can effectively target tumors, transport cargo, and facilitate the translocation of various payloads into the cytosol without needing cell receptor interactions or membrane pore formation.^20–25^ Previous studies have successfully harnessed pHLIP to deliver various drug molecules (including peptides) into solid tumors and metastatic sites in animal models.^26–31^

A promising payload for targeted tumor therapies involves antigenic peptides for presentation on the major histocompatibility complex (MHC). MHC molecules can present antigenic peptides to cytotoxic T lymphocytes, initiating an immune response.^32^ Specifically, antigenic peptides presented on MHC class I molecules can be recognized by cytotoxic CD8+ T cell receptors (TCRs), triggering the release of perforin and granzyme B, ultimately leading to target cell death.^33, 34^ In the context of cancer, cancer cells often display unique peptide-MHC complexes due to various alterations in their proteome, such as protein mutations, aberrant post-translational modifications, and other cellular processes.^35–37^ These unique peptides, called neoantigens, are absent in healthy cells and enable the immune system to selectively target and eliminate cancer cells with high efficiency, particularly in cancers with a high mutational burden.^38^ However, targeting neoantigens in cancers with a low mutational burden or heterogenous neoantigen expression has shown limited success.^39^ Additionally, the negative selection of self-reactive T cells can prevent neoantigens from having strong anti-cancer activity.^40^ Therefore, delivering highly antigenic peptides that are orthogonal to endogenous neoantigens and prompting their presentation on MHC molecules represents a potential strategy for enhancing treatment outcomes.

Prior work by Irvine and colleagues highlighted the benefit of delivering antigenic peptides in a non-specific manner by using cell-penetrating peptides. This approach led to the improved display of antigenic peptides on MHC and the development of T cells specifically targeting the desired epitope.^41^ An advantage of cytosolic delivery of antigenic peptides is that it bypasses endosomal processing of peptides, which can lead to significant degradation of MHC-binding peptide epitopes before they can be presented on MHC.^42^ Additionally, recent clinical trials have demonstrated the effectiveness of delivering cancer-specific neoantigens for display on MHC in use for cancer therapy.^43^

Building upon these findings, we posed that pHLIP could deliver antigenic peptides to cancer cells and this would offer a novel approach to selectively activate the immune system against cancer cells. Here, we showed that pHLIP conjugated to the model antigen SIINFEKL is selectively translocated through the membrane of cells in low pH environments and is subsequently displayed on MHC (Figure 1). The peptide epitope SIINFEKL (OVA) is derived from ovalbumin and there are no human equivalents, thus making it an orthogonal antigenic peptide with high affinity towards MHC molecules. Notably, we demonstrate that melanoma cells in acidic environments showed enhanced MHC display of the target epitope and increased recognition and activation by CD8+ T cells. These findings highlight the potential of pHLIP-mediated delivery of immunomodulatory agents as an anti-cancer therapy.

**Figure 1.**
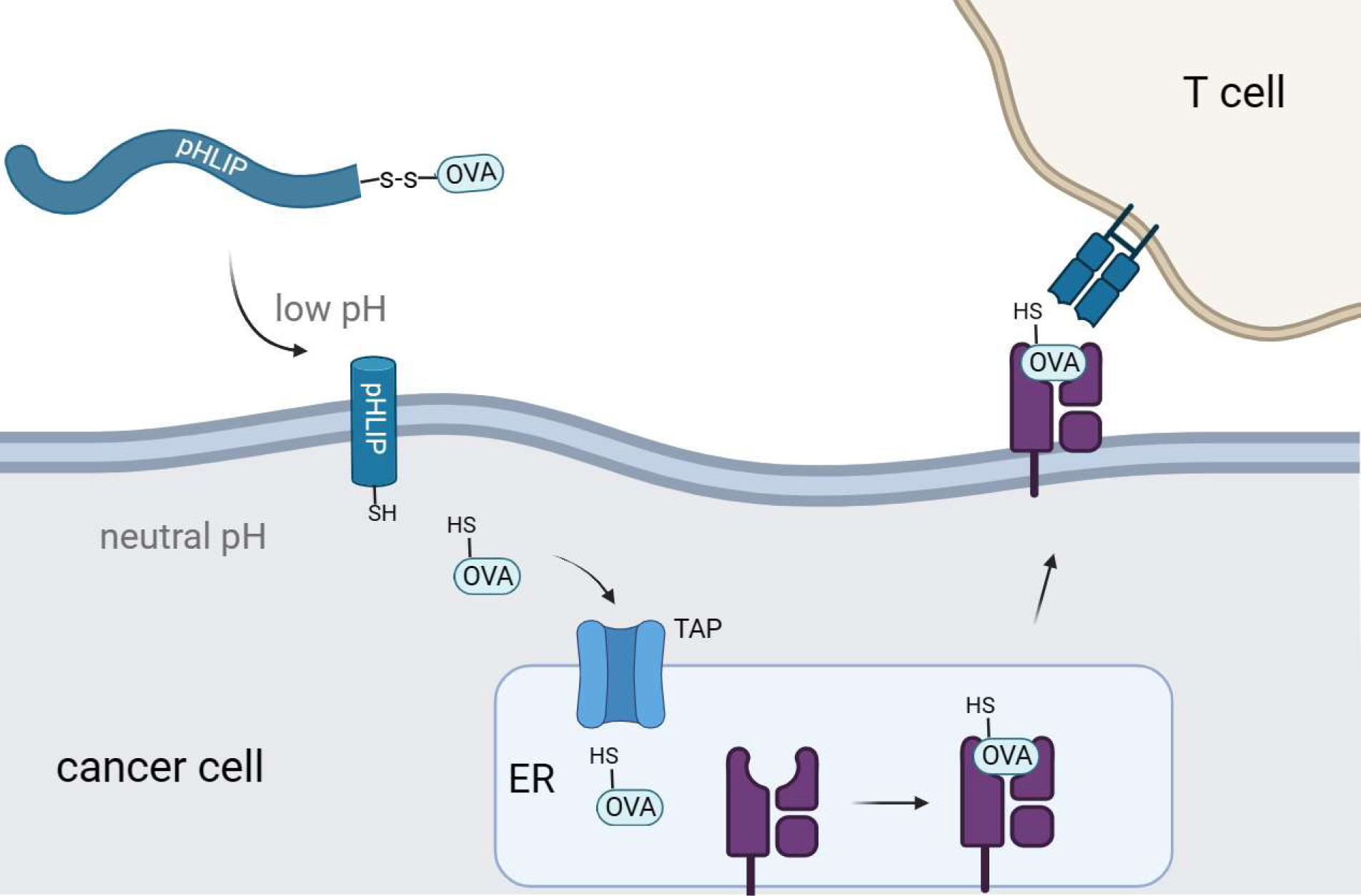
Schematic representation of pHLIP-mediated translocation of antigenic peptides for display on MHC. Intracellular delivery of the OVA peptide results in transportation to the endoplasmic reticulum for eventual display on MHC.

## Results and Discussion

We sought to link the OVA peptide to pHLIP *via* a chemical handle that would selectively uncouple upon arrival in the cytosolic space. While there are numerous strategies compatible with our system, we selected to connect OVA to pHLIP via a disulfide bond. The rationale behind employing disulfide conjugation was to facilitate the release of the peptide in the reducing environment of the cytosol to enable its entry into the antigen presentation pathway. To accomplish this strategy, it would be necessary to introduce a thiol group into the sequence of OVA. Our initial goal was to identify a site on OVA where the addition of a cysteine would have minimal impact on MHC binding and recognition by OVA-specific TCRs, considering that structural alterations on peptide sequences can significantly influence both parameters.^44, 45^

To assess changes in peptide affinity to MHC complexes from the introduction of cysteine to OVA, we used the RMA-S stabilization assay. The RMA-S cell line, which lacks the transporter associated with antigen processing (TAP), can be used to isolate the effect of peptide affinity because it lacks the ability to intracellularly process peptides for presentation.^46, 47^ Under low-temperature conditions (22-26°C), RMA-S cells present low-affinity pMHC complex on their cell surface. Increasing the temperature to 37°C causes low-affinity pMHC complex to dissociate, become internalized, and degraded. However, introducing a peptide with high affinity allows the pMHC complexes to remain stable at higher temperatures. The peptide-MHC binding affinity, specifically to the H-2K^b^ haplotype on RMA-S cells, is quantified using flow cytometry with a fluorescent anti-H-2K^b^ antibody.

Cysteine-containing OVA peptides were synthesized using a standard solid-phase peptide synthesis (SPPS) approach. Cysteine residues were introduced in sites within the OVA peptide that we projected would minimally impact binding to MHC molecules. These included the addition of cysteine to the termini of the sequence as well as the replacement of certain residues with cysteine (Figure S1). To analyze their affinity to MHC complexes, RMA-S cells were incubated with peptides at 26°C, allowing for the exchange of existing low-affinity pMHC complexes. Afterwards, the cells were then warmed to 37°C and treated with an APC-conjugated anti-mouse H-2K^b^ antibody. As expected, RMA-S cells treated with unmodified OVA peptide displayed high levels of MHC presentation on the cell surface (Figure 2A). The peptide SNFVSAGI (cntPEP) was used as a negative control as it has been reported to not appreciably bind to MHC.^45, 48^ Satisfyingly, the introduction of cysteine was well tolerated in most of the cysteine-modified OVA peptides; in particular, the cysteine introduction was better tolerated when the cysteine residues were located near the N-terminus. Nonetheless, all the cysteine-modified OVA peptides demonstrated substantial stabilization of the pMHC complex.

**Figure 2.**
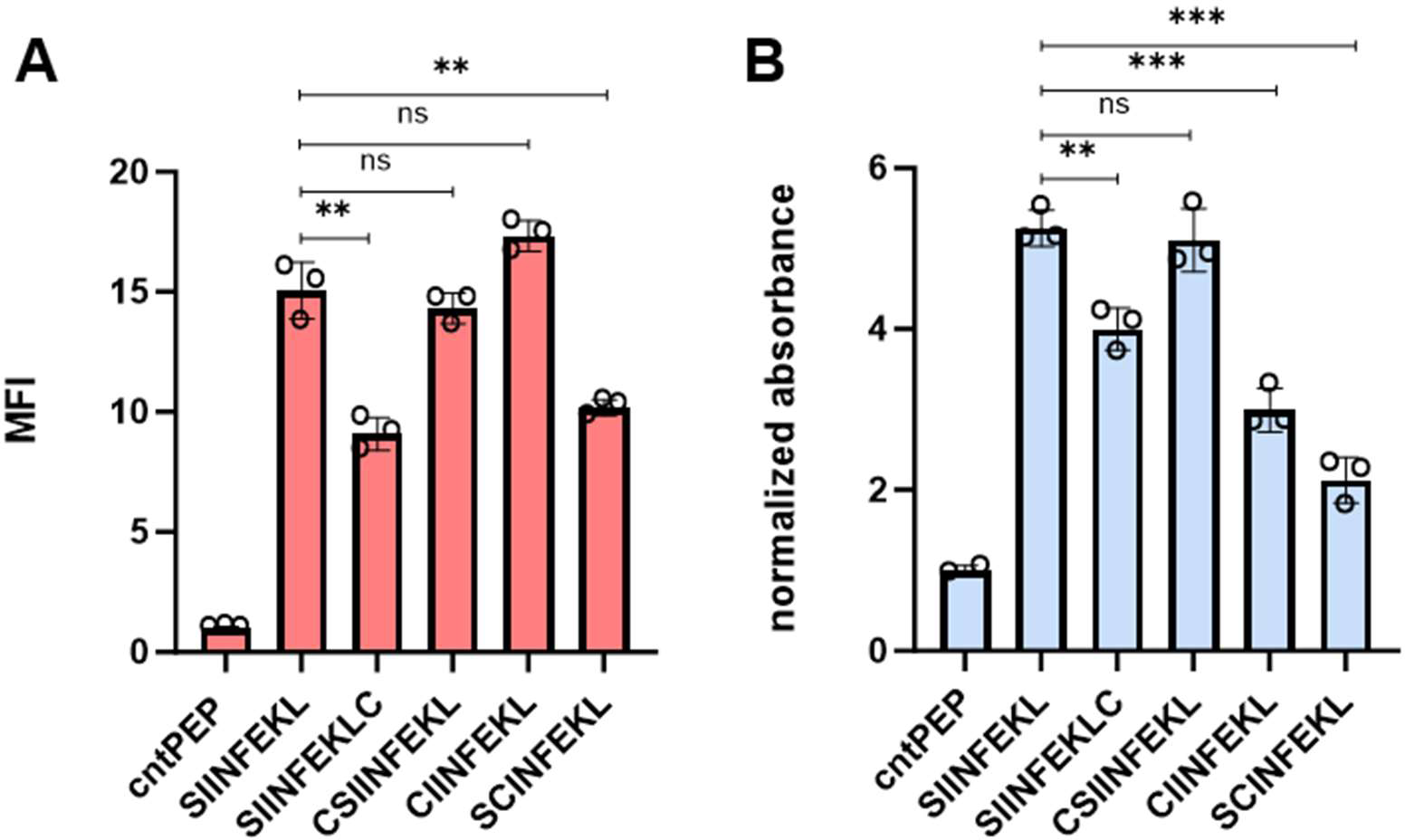
(A) Flow cytometry analysis of RMA-S cells. RMA-S cells were incubated with 20 μM peptide for 6 hours and pMHC complex display was quantified via APC conjugated anti-mouse H-2K^b^ antibodies. MFI for each group is the fold change in fluorescence intensity over the cntPEP. (B) RMA-S cells were incubated with 20 μM peptide and co-cultured with B3Z T cells for 8 hours at an effector-to-target ratio of 1:1. β-galactosidase expression was analyzed by measuring the hydrolysis of the colorimetric reagent CPRG on a plate reader at 570 nm and the data presented has been normalized to the absorbance from cntPEP. Data are represented as mean ± SD of biological replicates (n=3). P-values were determined by a two-tailed *t*-test (** p < 0.01, *** p < 0.001, ns = not significant).

Presentation of a cysteine-modified OVA peptide on MHC is critical for the success of our strategy, Previous findings, conducted by us and others, have indicated that changes to the OVA structure can potentially abolish TCR binding and subsequent T cell activation.^45,,49^. Additionally, we introduced cysteines residues in positions reported to have the least disruptive effects in order to minimize any unfavorable interactions with SIINFEKL-specific TCRs.^50^ We used an effector cell line, B3Z T cells, to evaluate T cell activation against target cells displaying cysteine modified OVA peptides. B3Z T cells have OVA-specific TCRs and express the enzyme β-galactosidase under the control of an IL-2 inducible promoter upon activation against target cells. The hydrolysis of chlorophenol red-β-D-galactopyranoside (CPRG) by β-galactosidase leads to a measurable color change, reflecting the levels of T cell activation. RMA-S cells were incubated with the peptides at 26°C before being co-cultured with B3Z T cells at 37°C. In this assay, β- galactosidase expression was quantified by measuring the hydrolysis of CPRG at 570 nm. All cysteine-containing OVA peptides demonstrated the capability to activate T cells to varying levels (Figure 2B). However, amongst the peptides tested, CSIINFEKL (CysOVA) exhibited both high MHC binding and TCR activation levels, comparable to those of the wild-type OVA peptide. Therefore, we proceeded with CysOVA for conjugation to pHLIP. Next, pHLIP was synthesized using SPPS with a corresponding cysteine added at its C-terminus. The disulfide conjugation between CysOVA and pHLIP was performed in solution, and the resulting conjugate (pHLIP-CysOVA) was purified via reverse phase high performance liquid chromatography (RP-HPLC).

To determine the secondary structures of pHLIP-OVA in the presence of a lipid bilayer at neutral and acidic pH, we utilized far-ultraviolet circular dichroism (CD) spectroscopy. pHLIP-OVA was incubated in the presence of 200 nM 1-palmitoyl-2-oleoyl-sn-glycero-3- phosphocholine (POPC) liposomes at pH 7.4 or pH 5.0. As shown in Figure 3A, a pH-dependent conformational shift from an unstructured random coil (pH 7.4) to an α-helix (pH 5.0) characteristic of pHLIP’s behavior was observed. Tryptophan (Trp) fluorescence spectroscopy was also employed to validate the pH-dependent membrane insertion propensity of pHLIP-OVA. Fluorescence emission from the two Trp residues in the sequence of pHLIP is sensitive to environment polarity and thus reports on lipid membrane insertion. When the pH was lowered from pH 7.4 to pH 5.0, we observed a λmax blue shift indicating the Trp residues transitioned to the hydrophobic environment of the lipid bilayer (Figure 3B). Together with the CD spectra, these results indicate that conjugation to CysOVA does not significantly impact the ability of pHLIP to insert across lipid membranes. This result agrees with the many previous studies showing that pHLIP can translocate a wide variety of C-terminally linked peptide cargoes across lipid bilayers.^20, 23, 25, 31, 51^

**Figure 3.**
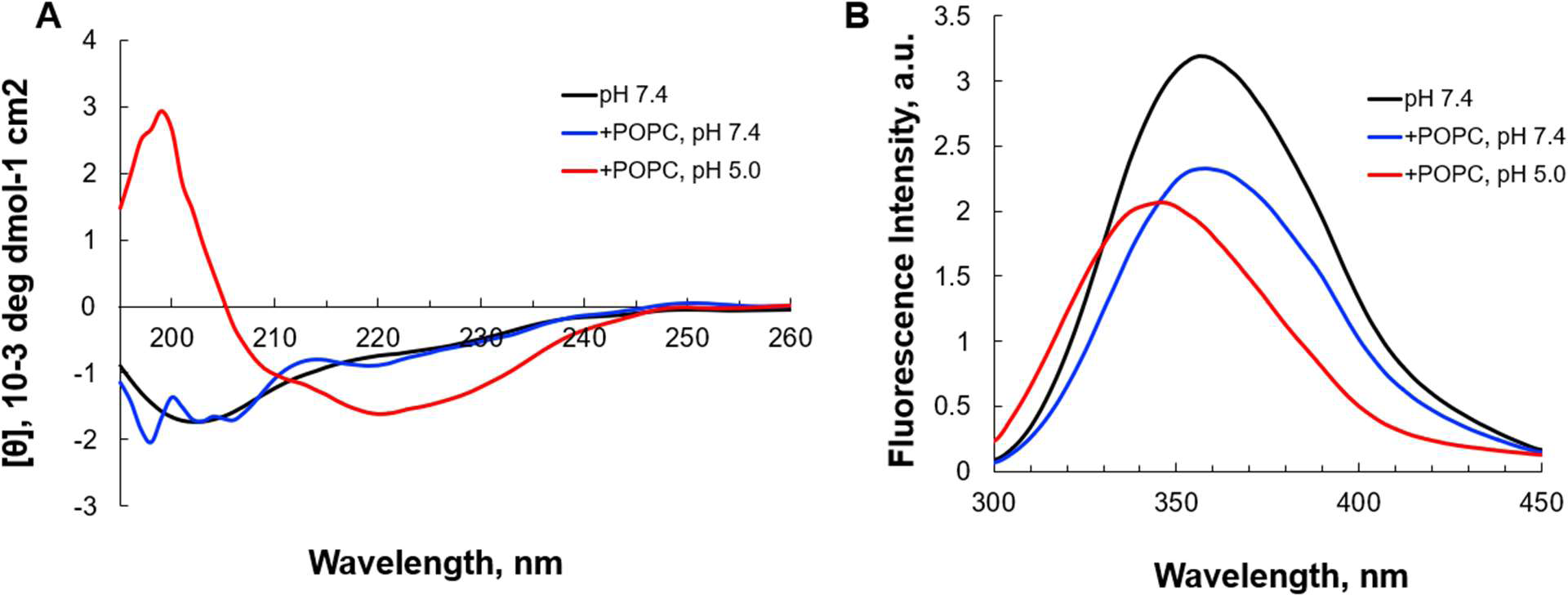
(A) Tryptophan fluorescence spectra of pHLIP-CysOVA (20 μM) in the presence of POPC vesicles at pH 7.4 or 5.0. (B) CD spectrum of pHLIP-CysOVA in the presence of POPC vesicles at pH 7.4 or 5.0.

To ensure that pHLIP-CysOVA cannot appreciably interact with MHC molecules before conversion into CysOVA, we determined the affinity of pHLIP-CysOVA to MHC before and after disulfide reduction. Given that the length of peptides that bind to MHC class I molecules are in the range of 8-12 amino acids, we anticipated that the disulfide bond would have to be uncoupled to generate CysOVA for improved MHC affinity. For this, we used the RMA-S stabilization assay to assess the relative affinity of the peptide to MHC before and after disulfide reduction at physiological pH. In this assay, RMA-S cells were treated with pHLIP-CysOVA in the presence or absence of β-mercaptoethanol (BME) to cleave the disulfide linkage to assess changes in MHC binding affinity. Our results showed that cleaving the disulfide bond to generate CysOVA stabilized the peptide-MHC complex on RMA-S cells and significantly improved antigen presentation on the cell surface (Figure 4A). Overall, these results indicate that pHLIP-CysOVA must be reduced into CysOVA for optimal display on MHC.

**Figure 4.**
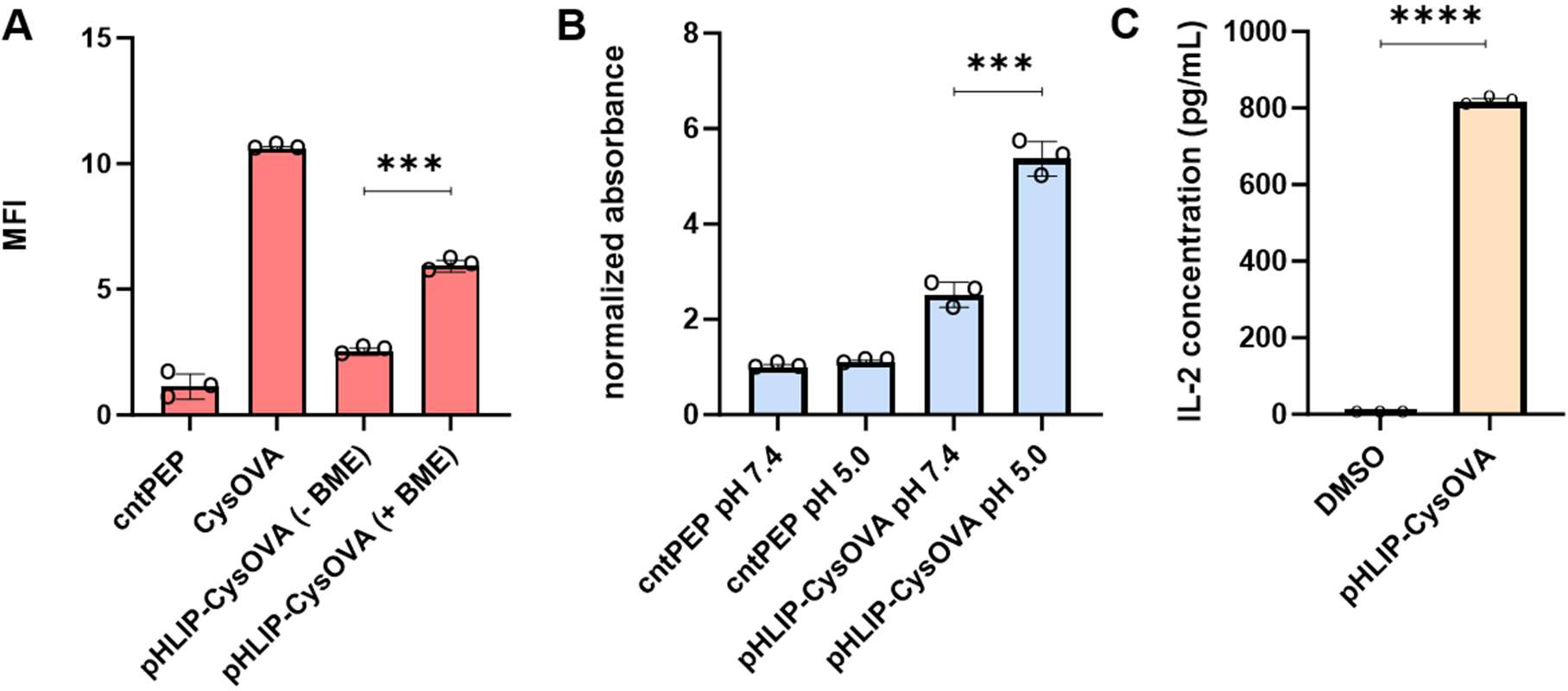
(A) RMA-S cells at physiological pH were incubated with 2.5 μM pHLIP-CysOVA for 6 hours in the presence or absence of BME. Cells were then labeled with anti-mouse H-2K^b^ antibody and analyzed via flow cytometry. MFI is mean fluorescence intensity of the level of fluorescence relative to the control peptide (B) 2.5 μM pHLIP-CysOVA was incubated with B16 cells for 5 mins before adjusting to the indicated pH for 10 mins. Cells were washed and co-cultured with B3Z T cells for 8 hours before lysing and measuring β-galactosidase activity via the colorimetric reagent CPRG on a plate reader at 570 nm and the data presented has been normalized to the absorbance from cntPEP. (C) B16 cells were treated with pHLIP-CysOVA for 5 mins before adjusting to pH 5 for 10 mins.

Subsequently, these cells were washed and co-cultured with B3Z T cells overnight, with secreted IL-2 levels measured through ELISA. Data are represented as mean ± SD of biological replicates (n= 3). P-values were determined by a two-tailed *t*-test (*** p < 0.001, **** p < 0.0001).

To evaluate the efficacy of pHLIP-CysOVA in enabling antigen presentation on MHC in cells within acidic microenvironments in TAP competent cells, DC2.4 dendritic cells were subjected to incubation with pHLIP-CysOVA under neutral and acidic conditions. We expect that treatment of pHLIP-CysOVA in acidic environments would improve cytosolic delivery of CysOVA and lead to better presentation on MHC. Cells were treated with pHLIP-CysOVA at either pH 7.4 or 5.0 for 10 mins at 37°C. Subsequently, cells were washed and incubated with APC-conjugated 25-D1.16 antibodies (specific to SIINFEKL bound to H-2K^b^) to quantify antigen presentation on MHC. Our data demonstrated a significant increase in antigen presentation on MHC in cells treated at low pH, indicating successful translocation of pHLIP-CysOVA across the membrane and entry into the antigen presentation pathway (Figure S2).

Finally, to demonstrate that treatment at low pH enables selective immune activation of pHLIP-CysOVA in a clinically relevant model, B16 melanoma cells were employed. B16 cells were incubated with pHLIP-CysOVA at either physiological or low pH for 10 min. Following this incubation, the cells were washed and co-cultured with effector cells (B3Z T cells). Once again, treatment at lower pH conditions significantly enhanced T cell activation against the target B16 cells (Figure 4B). Additionally, ELISA results confirmed that pHLIP-CysOVA treatment at low pH-induced T cell activation, as there was elevated IL-2 cytokine secretion from B3Z T cells (Figures 4C). However, cells treated with CysOVA peptide alone did not show any pH dependence in terms of T cell activation (Figure S3). Taken together, these findings demonstrate selective activation of T cells in response to pHLIP-CysOVA treatment within acidic environments.

## Conclusion

In this study, we have described a targeted approach to deliver antigenic peptides specifically to cancer cells, thereby aiding their presentation on MHC molecules to enhance immune activation. Our method used pHLIP to deliver CysOVA directly to the cytosol, eliminating the requirements for cross-presentation.^52^ By avoiding this step, we prevent significant peptide degradation that typically occurs within the acidic endosomal environment before translocation into the cytosol, ensuring efficient presentation of the peptide on MHC molecules.^53^

A key advantage of our approach is the versatility of pHLIP, which has previously demonstrated the ability to translocate a wide range of molecules, including hydrophilic peptides, across cell membranes.^25^ We envision that our strategy can be adapted to deliver various potential neoantigens, thereby broadening its applicability and therapeutic potential. Additionally, it was recently reported that MHC peptides tagged with distinct covalent small-molecule inhibitors can be targeted by immune cells for immunotherapy applications.^54^ Consequently, we anticipate that pHLIP can serve as a delivery vehicle for chemically modified peptides, potentially used alongside antibodies or CAR-T cells to enable precision immunotherapy against cancer.^55^

## Acknowledgement

This study was supported by the NIH grant R21 AI172180.

## Supporting Information

Additional figures, tables, and materials/methods are included in the supporting information file.

## Notes

### Competing Interest Statement

The authors have declared no competing interest.

