## Supporting Information for "Targeted Acidosis Mediated Delivery of Antigenic MHC-Binding Peptides"

Joey J. Kelly[1], Emily Akron[2], Damien Thévenin[2], and Marcos M. Pires[1*]

[1]Department of Chemistry
University of Virginia
Charlottesville, VA, United States

[2]Department of Chemistry
Lehigh University
Bethlehem, PA, United States

### Table of Contents


| | |
|---|---|
| **Supporting Figures** | **S3-S5** |
| Figure S1. Chemical structures of cysteine-containing OVA peptides | S3 |
| Figure S2. Flow cytometry analysis of pHLIP-CysOVA treated DC2.4 cells with 25-D1.16 antibody | S4 |
| Figure S3. T cell activation of pHLIP-CysOVA Treated B16 cells | S5 |
| **Materials and Methods** | **S6-S8** |
| Materials | S6 |
| Mammalian cell culture | S6 |
| RMA-S Stabilization Assay | S6 |
| B3Z T cell activation | S6 |
| Sample Preparation for CD and Tryptophan Emission Spectroscopy | S7 |
| CD Spectroscopy | S7 |
| B3Z Targeting pHLIP-OVA Treated B16 Cells | S7 |
| 25-D1.16 Labeled pHLIP-CysOVA Treated DC2.4 Cells | S7 |
| B3Z IL-2 Secretion | S8 |
| **Synthesis and Characterization** | **S9-S21** |
| Scheme S1: SIINFEKL | S9 |
| Scheme S2: pHLIP | S11 |
| Scheme S3: pHLIP-OVA | S12 |
| Scheme S4: SCINFEKL | S13 |
| Scheme S5: CIINFEKL | S15 |
| Scheme S6. CSIINFEKL | S17 |
| Scheme S7: SIINFEKLC | S19 |
| Scheme S8: SNFVSAGI | S21 |

#### SUPPORTING FIGURES

| cntPEP | SIINFEKL |
|---|---|
| 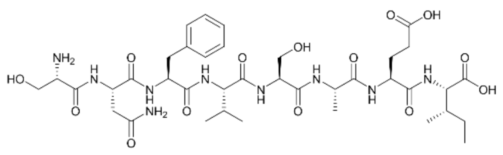 | 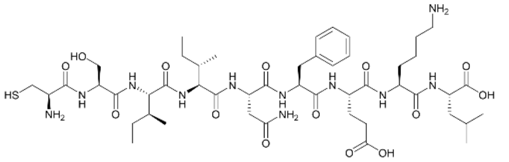 |
| **SIINFEKLC** | **CIINFEKL** |
| 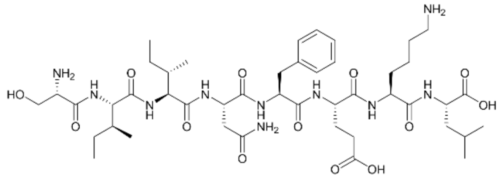 | 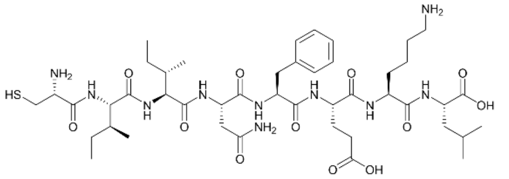 |
| **CSIINFEKL** | **SCINFEKL** |
| 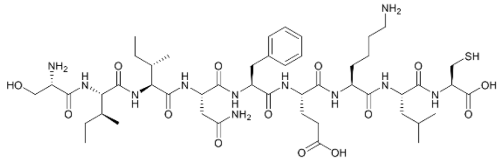 | 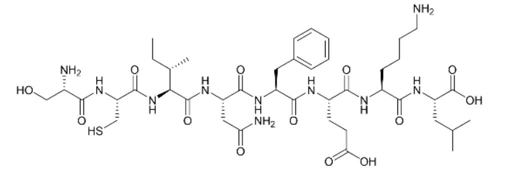 |

**Figures S1**. Chemical structure of cystine-containing OVA peptides.

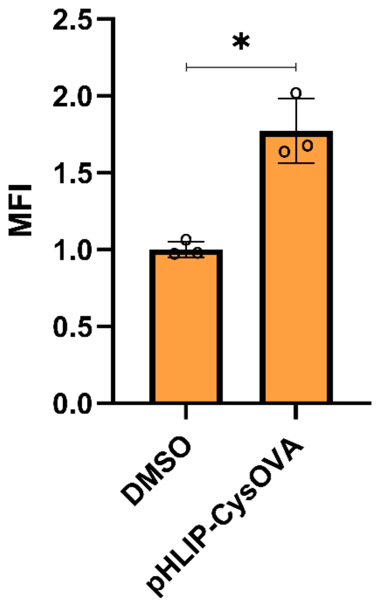


**Figures S2**. DC2.4 cells were incubated with peptide for 5 mins before adjusting to pH 5 for 10 mins. Cells were washed and incubated with APC labeled 25-D1.16 antibody (specific for SIINFEKL on H-2K$^b$) for 1 h and analyzed via flow cytometry. Data are represented as mean ± SD of biological replicates (n= 3). P-values were determined by a two-tailed *t*-test (* $p < 0.05$).

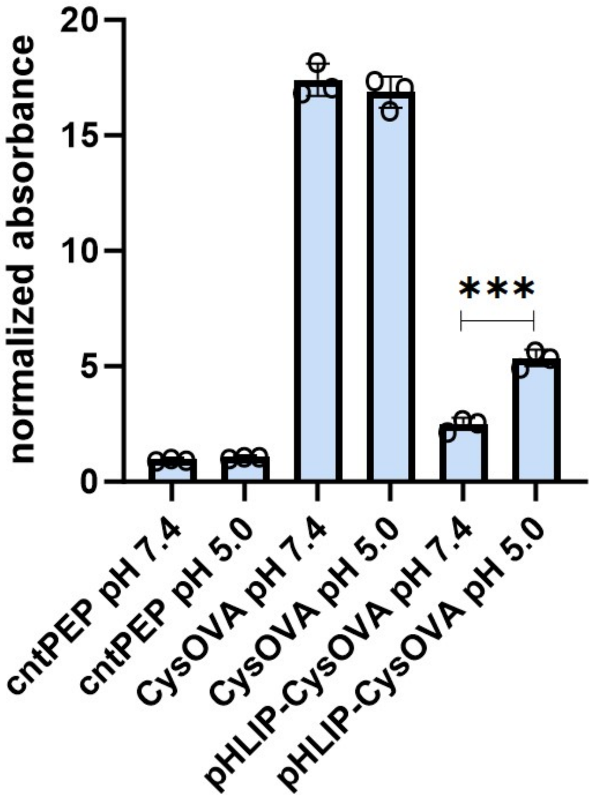


**Figures S3**. 2.5 μM pHLIP-CysOVA was incubated with B16 cells for 5 mins before adjusting to the indicated pH for 10 mins. Cells were washed and co-cultured with B3Z T cells for 8 hours before lysing and measuring β-galactosidase activity via the colorimetric reagent CPRG on a plate reader at 570 nm. Data are represented as mean ± SD of biological replicates (n= 3). P-values were determined by a two-tailed *t*-test (*** $p < 0.001$).

**Materials.**

All peptide related reagents and protected amino acids were purchased from Chem-Impex. APC-labeled anti-mouse H-2K$^d$/H-2D$^d$ antibody was purchased from Biolegend._Pooled Human Serum was purchased from Sigma Aldrich. Dulbecco's Modified Eagle's Medium (DMEM) and Roswell Park Memorial Institute (RPMI) 1640 medium were purchased from VWR. Fetal Bovine Serum (FBS) was purchased from R&D Systems. Penicillin-streptomycin and mouse IL-2 ELISA kits was purchased from Sigma-Aldrich. All other organic chemical reagents were purchased from Fisher Scientific or Sigma Aldrich and used without further purification. All compounds are >95% pure by HPLC analysis.

**Experimental Methods.**

**Mammalian Cell Culture.** RMA-S cells were a kind gift from Dr. John Sampson. RMA-S cells were maintained in RPMI 1640 media supplemented with 10% fetal bovine serum, 50 IU/mL penicillin, 50 ug/mL streptomycin,1X a MEM non-essential amino acid solution (ThermoFisher) and cultured in a humidified atmosphere of 5% $CO_2$ at 37$^o$C. B3Z cells were kindly provided by Dr. Aaron Esser-Kahn and maintained in RPMI 1640 media supplemented with 10% fetal bovine serum, 50 IU/mL penicillin, 50 ug/mL streptomycin and cultured in a humidified atmosphere of 5% $CO_2$ at 37$^o$C. B16 cells were kindly provided by Dr. Victor Engelhard and also maintained in PRMI 1640 media supplemented with 10% fetal bovine serum, 50 IU/mL penicillin, 50 ug/mL streptomycin and cultured in a humidified atmosphere of 5% $CO_2$ at 37$^o$C.

**RMA-S Stabilization Assay**. $10^5$ RMA-S cells were seeded in a treated 96 well plate at 37$^o$C overnight. The next day, RMA-S cells were moved to a 26$^o$C incubator for 24-48 hours. Following the incubation period, cells were incubated with 20 $\mu$M peptides in culture media at indicated concentrations for 1 hour at 26$^o$C before being moved to the 37$^o$C incubator for 6 hours. For assays involving $\beta$-mercaptoethanol, 100 $\mu$M of BME was added along with peptide during these incubation periods. The media was then replaced with a 1:100 dilution of APC-labeled anti-mouse H-2K$^d$/H-2D$^d$ antibody in culture media for 1 hour at 4$^o$C. Cells were removed from the well plate by vigorous pipetting, fixed with 2% formaldehyde solution, and analyzed using the Attune NxT Flow Cytometer (Thermo Fischer) equipped with a 637 nm laser with 670/14 nm bandpass filter.

**B3Z T Cell Activation**. $10^5$ RMA-S cells were seeded in a treated 96 well plate at 37$^o$C overnight. The following day the culture media was replaced with media containing 20 $\mu$M peptide along with $10^5$ B3Z cells in culture media and co-incubated overnight. Cells were then spun down at 500xg for 5 mins and washed with 1X PBS a total of two times. Lysis buffer containing 0.02% saponin, 500 $\mu$M CPRG reagent, 100 mM $MgCl_2$, and 100 mM $\beta$-mercaptoethanol in 1X PBS was added to each well. After 2-4 hours absorbance 570 was recorded using a BioTek Epoch 2 microplate reader.

#### Sample Preparation for CD and Tryptophan Emission Spectroscopy

pHLIP-OVA was solubilized to 20 μM in 5 mM sodium phosphate buffer (pH 8.0). The peptide was diluted to a final concentration of 5 μM before analysis. Liposomes were prepared by drying 1-palmitoyl-2-oleoyl-sn-glycero-3-phosphocholine (POPC) as a thin film and desiccation under vacuum for at least 24 h. Lipids were rehydrated in 5 mM sodium phosphate (pH 8.0) for at least 30 min at 37°C with periodic gentle vortexing. The resulting large multilamellar vesicles were rapidly frozen and thawed seven times and subsequently extruded through a polycarbonate membrane with 200 nm pores using a Mini-Extruder (Avanti Polar Lipids) to produce large unilamellar vesicles (LUVs). pHLIP-OVA was incubated with the resulting LUVs at a 1:300 ratio. pH was adjusted to the desired experimental values with HCl and the samples were incubated at RT for 30 min prior to analysis. Fluorescence emission spectra were recorded using a Fluorolog-3 spectrofluorometer (HORIBA). The excitation wavelength was set at 280 nm, and the emission spectra were measured from 300 to 450 nm. The excitation and emission slit widths were set to 5 nm.

#### CD Spectroscopy

Far-ultraviolet CD spectra were recorded on a Jasco J-815 CD spectrometer equipped with a Peltier thermally controlled cuvette holder (Jasco). Samples were measured in 0.1 cm quartz cuvette. CD intensity was expressed as mean residue molar ellipticity [$\theta$] calculated by the following equation:

$[\theta] = \theta_{obs}/10lcn$ (in deg $cm^{-2}$ $dmol^{-1}$)

$\theta_{obs}$ is the observed ellipticity in millidegrees, $l$ is the optical path length in centimeters, $c$ is the molar peptide concentration, and $n$ is the number of amino acid residues. Spectra were recorded from 260 to 200 nm in 1 nm intervals at a 100 nm/min scan rate. Five scans were averaged for each sample. The spectrum of POPC liposomes was subtracted out from samples containing pHLIP-OVA in the presence of POPC.

**B3Z Targeting pHLIP-OVA Treated B16 Cells.** B16 cells were seeded in a treated 96-well plate at 37$^{o}$C overnight. The following day cell media was removed and treatment solution with 2.5 μM of pHLIP-OVA and incubated at 37$^{o}$C for 5-10 min. The pH was then adjusted to a pH of 5 by using a pre-established volume of DMEM, pH 2.0 buffered with citric acid, and the plate was incubated for 10 min. After the treatment, the media was washed once with DMEM and $10^{5}$ B3Z cells were added to each well. B3Z and B16 cells were co-cultured for 8 hours before being spun down at 500xg for 5 min and washed with 1X PBS a total of two times. Lysis buffer containing 0.02% saponin, 500 mM CPRG reagent, 100 mM $MgCl_2$, and 100 mM β-mercaptoethanol in 1X PBS was added to each well. After 2-4 hours absorbance 570 was recorded using a BioTek Epoch 2 microplate reader

#### 25-D1.16 Labeled pHLIP-CysOVA treated DC2.4 Cells

DC2.4 cells were seeded in a treated 96-well plate at 37$^{o}$C overnight. The following day cell media was removed and treatment solution with indicated concentration of pHLIP-OVA and incubated at 37$^{o}$C for 5-10 min. The pH was then adjusted to the desired value using a pre-established volume of DMEM, pH 2.0 buffered with citric acid, and the plate was incubated for 10 min. The media was replaced with a 1:100 dilution of the APC conjugated SIINFEKL-H-2K$^{b}$ specific antibody clone 25-D1.16 for 2 hours at 4$^{o}$C. Cells were fixed with 2% formaldehyde solution and analyzed using the Attune NxT Flow Cytometer (Thermo Fischer) equipped with a 637 nm laser with 670/14 nm bandpass filter.

**B3Z IL-2 Secretion**

B16 cells were seeded in a treated 24-well plate at 37$^{o}$C overnight. The following day cell media was removed and treatment solution with 2.5 μM of pHLIP-OVA and incubated at 37$^{o}$C for 5-10 min. The pH was then adjusted to a pH of 5 by using a pre-established volume of DMEM, pH 2.0 buffered with citric acid, and the plate was incubated for 10 min. After the treatment, the media was washed once with DMEM and $10^{6}$ B3Z cells were added to each well. B3Z and B16 cells were co-cultured overnight. IL-2 released by B3Z or B16 cells in cell supernatants were detected by IL-2 ELISA kits according to the manufacture's instruction.

**Scheme S1. Synthesis of SIINFKEL**

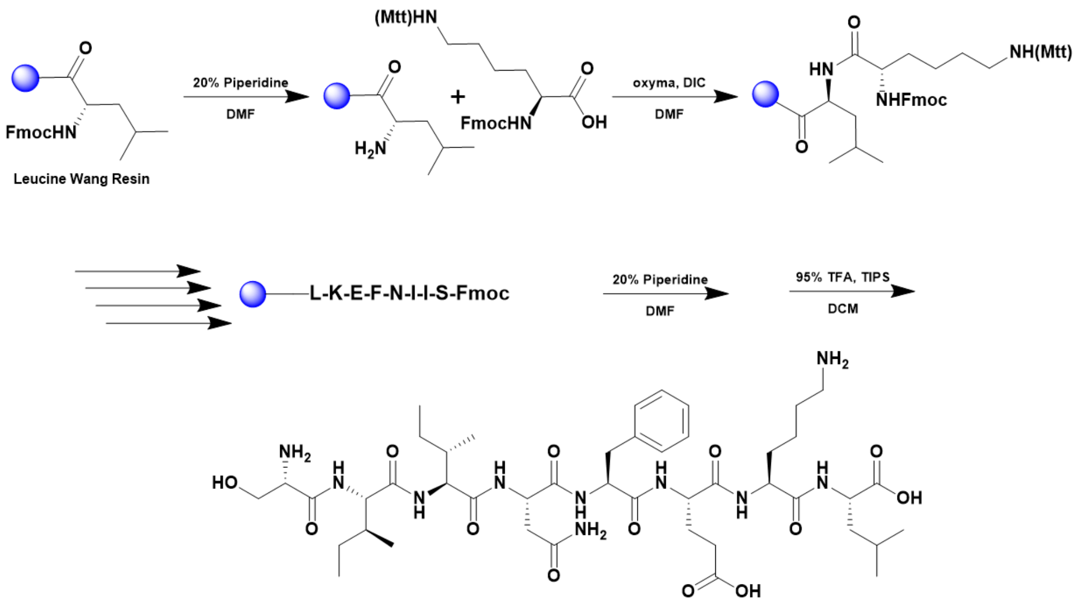


A 25 mL vessel of CEM discover bio manual peptide synthesizer was charged with 0.25 mmol of leucine wang resin. The Fmoc group was removed by using a 20% piperidine solution in DMF (10 mL). Using Synergy software, the deprotection protocol was run. The piperidine solution was drained and the resin was washed with DMF (4 x 10 mL). Fmoc-L-lysine(Boc)-OH (5 eq, 1.25 mM) along with Oxyma (5 eq, 1.25 mM) and DIC (5 eq, 1.35 mmol) in DMF was added to the reaction vessel and the coupling protocol was run. The amino acid solution was drained, and the resin was washed with DMF (2 x 10 mL). The fmoc removal and coupling procedure was repeated as before using the same equivalencies for the remaining amino acids. To remove the peptide from resin, a TFA cocktail solution (95% TFA, 2.5% TIPS, and 2.5% DCM) was added to the resin and agitated for 2 hours. The resin was filtered, and the resulting solution was concentrated in vacuo. The peptide was trituated with cold diethyl ether and purified using reverse phase HPLC using $H_2O/CH_3CN$. The sample was analyzed for purity using a Waters 1525 Binary HPLC Pump using a Phenomenex Luna 5u C8(2) 100A (250 x 4.60 mm) column; gradient eluted with $H_20/CH_3CN$. Molecular weight was confirmed using high resolution electrospray ionization mass spectrometry (HRMS, ESI/MS) analyses obtained on an Agilent 6545B Q-TOF LC/MS equipped with 1260 infinity II LC system with auto sampler. The final peptide product was lyophilized and stored at -20$^o$C until further use.

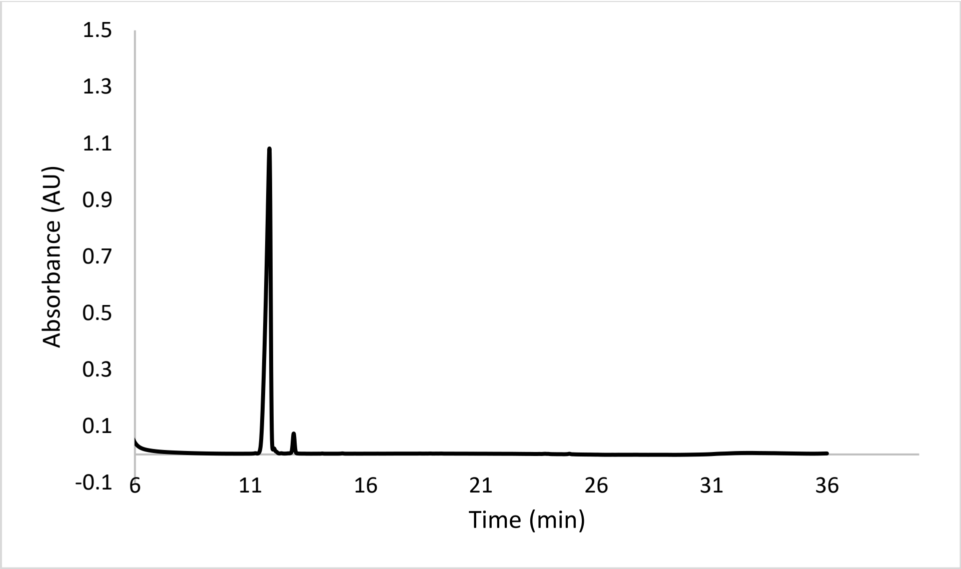


ESI-MS calculated [M+H$^+$]: 963.5515, found 963.5506.

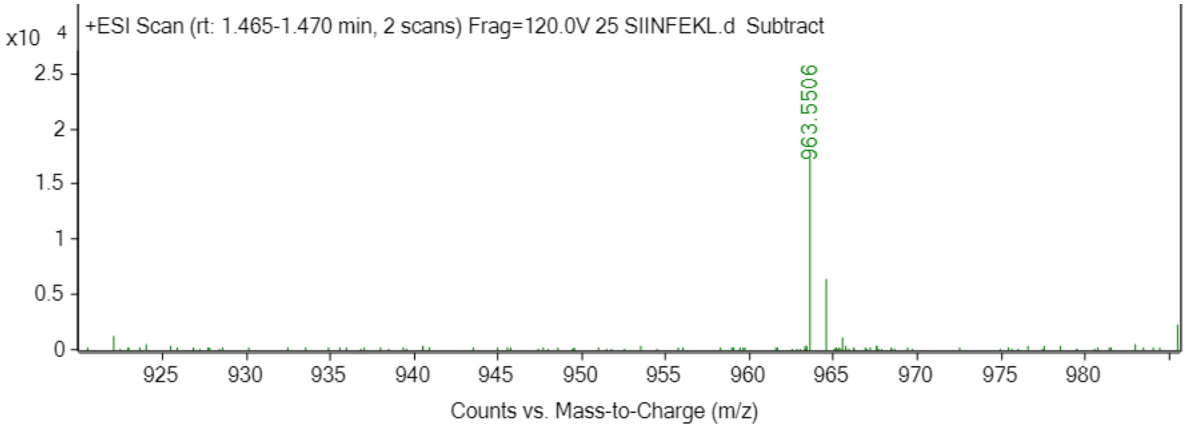

**Scheme S2. Synthesis of pHLIP**

pHLIP (GGEQNPIYWARYADWLFTTPLLLLDLALLVDADEGTCG) was prepared by Fmoc solid-phase peptide synthesis using rink amide resin (CEM, 0.19 mmol/g loading capacity) on a CEM Liberty Blue™ microwave peptide synthesizer. Each amino acid solubilized at 0.2 M in DMF was coupled using DIC or HBTU as the activator and oxyma or DIEA as the activator base. After each coupling step, Fmoc protecting groups were removed with 6% piperazine and 0.1 M HOBt in DMF. The peptide was cleaved from resin using a cocktail of trufluoroacetic acid (TFA)/phenol/water/thioanisole/1,2-ethanodithiol (82.5/5/5/5/2.5, v/v) for 2 h at room temperature. The solution was filtered to remove the resin and concentrated prior to precipitation with ice cold diethyl ether. pHLIP was purified by reverse-phase high-performance liquid chromatography (RP-HPLC) on a Phenomenex Luna Omega, 5μm, 250 mm × 21.2 mm C18 column; flow rate of 5 mL/min; phase A being water with 0.1% TFA; phase B being acetonitrile with 0.1% TFA; 60 min gradient from 95:5 to 0:100 A:B). Peptide identity was confirmed by matrix-assisted laser desorption ionization time-of-flight (MALDI-TOF) mass spectrometry (Shimadzu 8020).

MALDI-TOF calculated [M+H$^+$]:4210.72, found 4209.81.

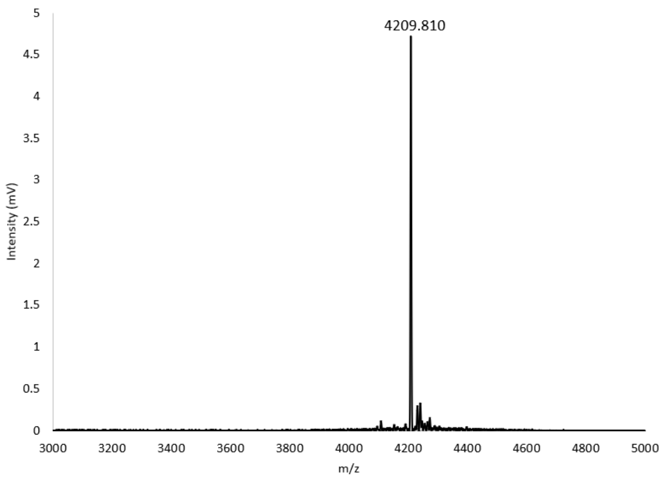

**Scheme S3. Synthesis of pHLIP-CysOVA**

The pHLIP peptide was dissolved to 1 mM in DMSO followed by the addition of 1.5 mM CysOVA in DMSO. The samples were thoroughly mixed before adding 1 M tris buffer pH 8.0 for 4 h. The resulting pHLIP-CysOVA was purified using a Waters 1525 Binary HPLC Pump using a Phenomenex Luna 5u C8(2) 100A (250 x 4.60 mm) column; gradient eluted with $H_20/CH_3CN$. Molecular weight was confirmed via a Shimadzu MALDI-TOF Mass Spectrometer (MALDI-8020).

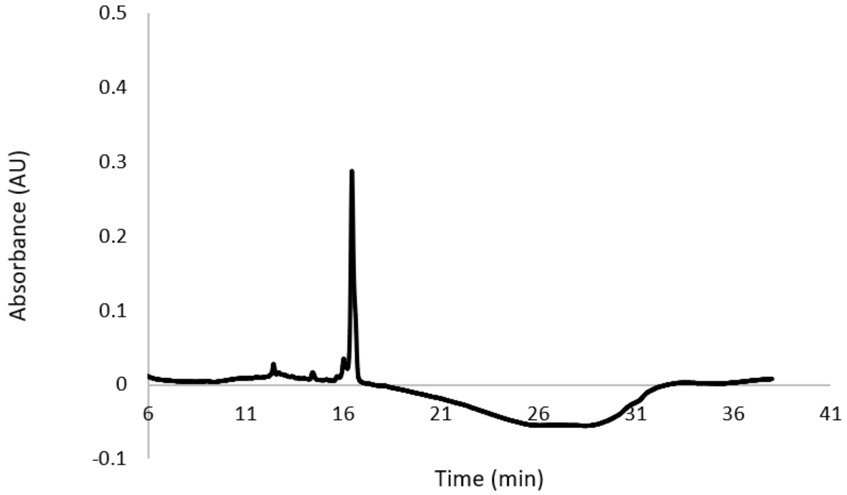


MALDI-TOF calculated [M+H$^+$]: 5277, found 5280.

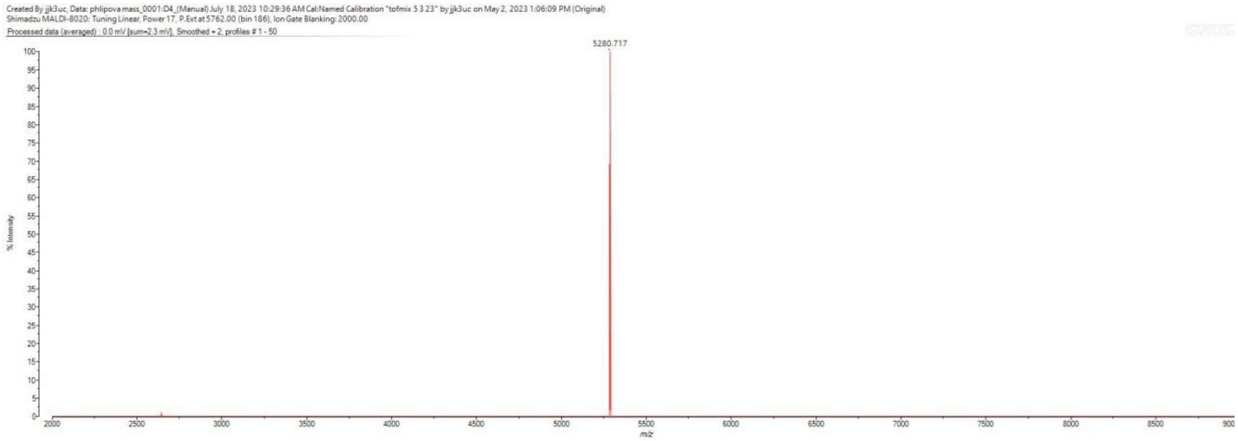

**Scheme S4. Synthesis SCINFEKL**

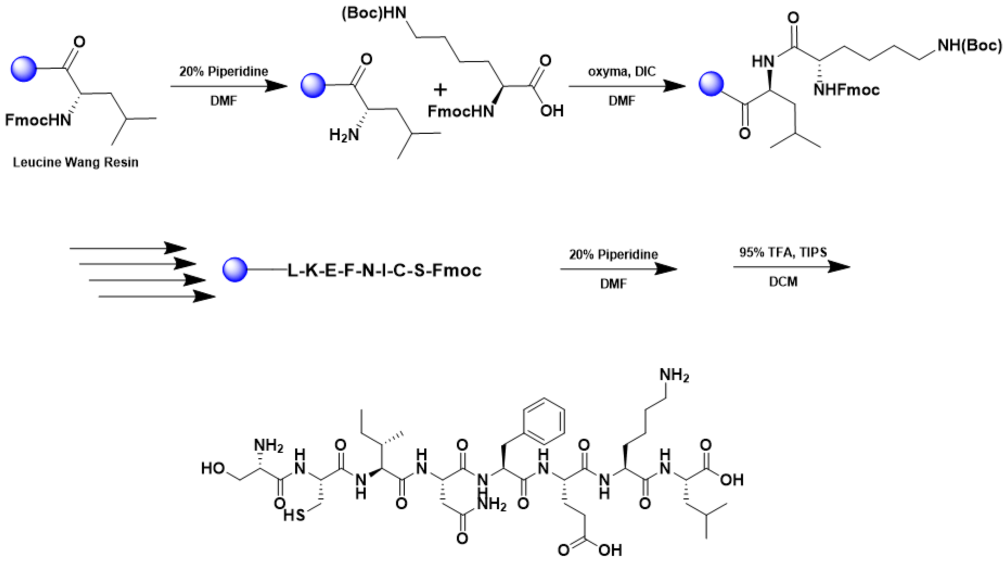


A 25 mL vessel of CEM discover bio manual peptide synthesizer was charged with 0.25 mmol of leucine wang resin. The Fmoc group was removed by using a 20% piperidine solution in DMF (10 mL). Using Synergy software, the deprotection protocol was run. The piperidine solution was drained and the resin was washed with DMF (4 x 10 mL). Fmoc-L-lysine(Boc)-OH (5 eq, 1.25 mM) along with Oxyma (5 eq, 1.25 mM) and DIC (5 eq, 1.35 mmol) in DMF was added to the reaction vessel and the coupling protocol was run. The amino acid solution was drained, and the resin was washed with DMF (2 x 10 mL). The fmoc removal and coupling procedure was repeated as before using the same equivalencies for the remaining amino acids. To remove the peptide from resin, a TFA cocktail solution (95% TFA, 2.5% TIPS, and 2.5% DCM) was added to the resin and agitated for 2 hours. The resin was filtered, and the resulting solution was concentrated in vacuo. The peptide was triturated with cold diethyl ether and purified using reverse phase HPLC using $H_2O/CH_3CN$. The sample was analyzed for purity using a Waters 1525 Binary HPLC Pump using a Phenomenex Luna 5u C8(2) 100A (250 x 4.60 mm) column; gradient eluted with $H_20/CH_3CN$. Molecular weight was confirmed using high resolution electrospray ionization mass spectrometry (HRMS, ESI/MS) analyses obtained on an Agilent 6545B Q-TOF LC/MS equipped with 1260 infinity II LC system with auto sampler. The final peptide product was lyophilized and stored at -20$^o$C until further use.

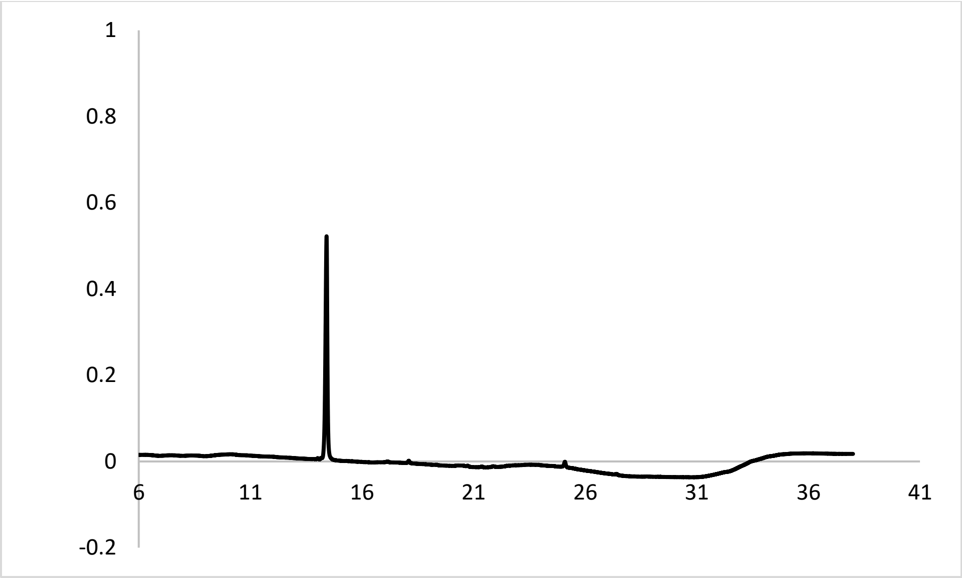


ESI-MS calculated [M+H$^+$]: 953.4761, found 953.4733.

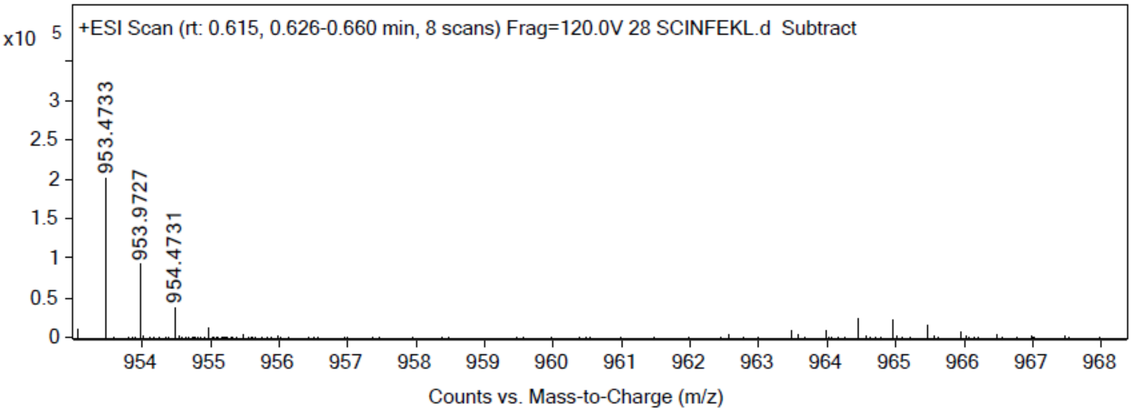

**Scheme S5. Synthesis CIINFKEL**

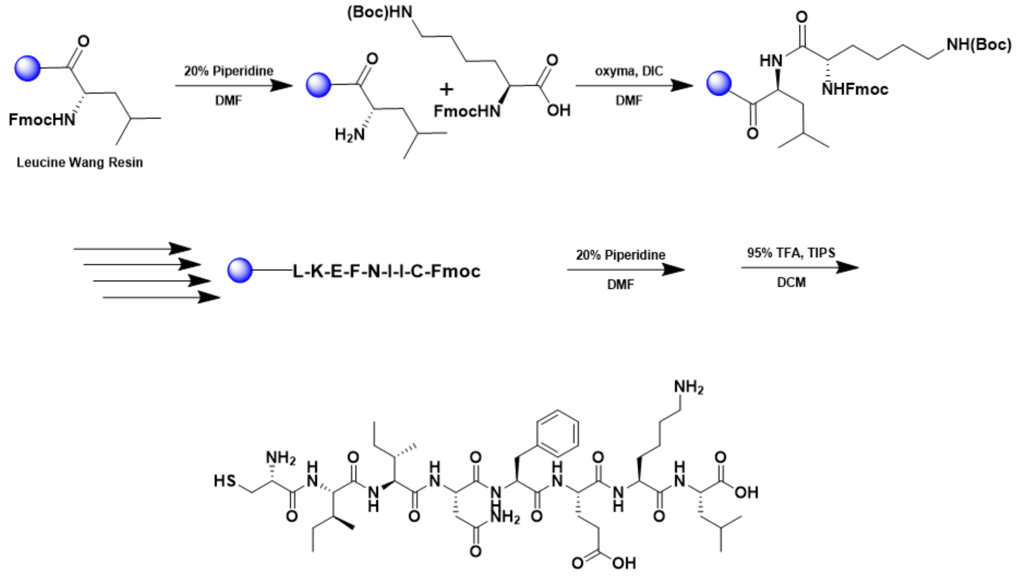


A 25 mL vessel of CEM discover bio manual peptide synthesizer was charged with 0.25 mmol of leucine wang resin. The Fmoc group was removed by using a 20% piperidine solution in DMF (10 mL). Using Synergy software, the deprotection protocol was run. The piperidine solution was drained and the resin was washed with DMF (4 x 10 mL). Fmoc-L-lysine(Boc)-OH (5 eq, 1.25 mM) along with Oxyma (5 eq, 1.25 mM) and DIC (5 eq, 1.35 mmol) in DMF was added to the reaction vessel and the coupling protocol was run. The amino acid solution was drained, and the resin was washed with DMF (2 x 10 mL). The fmoc removal and coupling procedure was repeated as before using the same equivalencies for the remaining amino acids. To remove the peptide from resin, a TFA cocktail solution (95% TFA, 2.5% TIPS, and 2.5% DCM) was added to the resin and agitated for 2 hours. The resin was filtered, and the resulting solution was concentrated in vacuo. The peptide was triturated with cold diethyl ether and purified using reverse phase HPLC using $H_2O/CH_3CN$. The sample was analyzed for purity using a Waters 1525 Binary HPLC Pump using a Phenomenex Luna 5u C8(2) 100A (250 x 4.60 mm) column; gradient eluted with $H_20/CH_3CN$. Molecular weight was confirmed using high resolution electrospray ionization mass spectrometry (HRMS, ESI/MS) analyses obtained on an Agilent 6545B Q-TOF LC/MS equipped with 1260 infinity II LC system with auto sampler. The final peptide product was lyophilized and stored at -20$^{o}$C until further use.

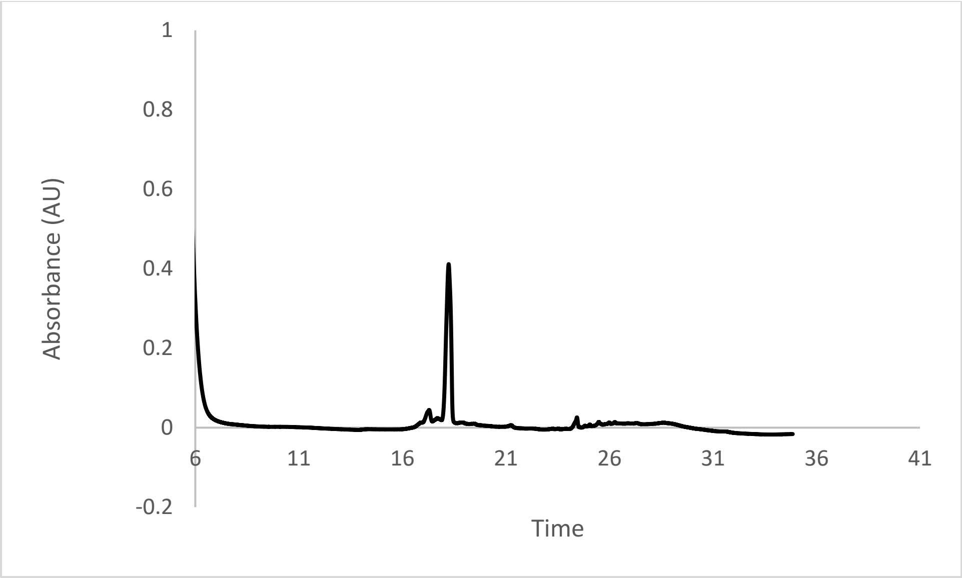


ESI-MS calculated [M+H$^+$]: 979.5258, found 979.5241.

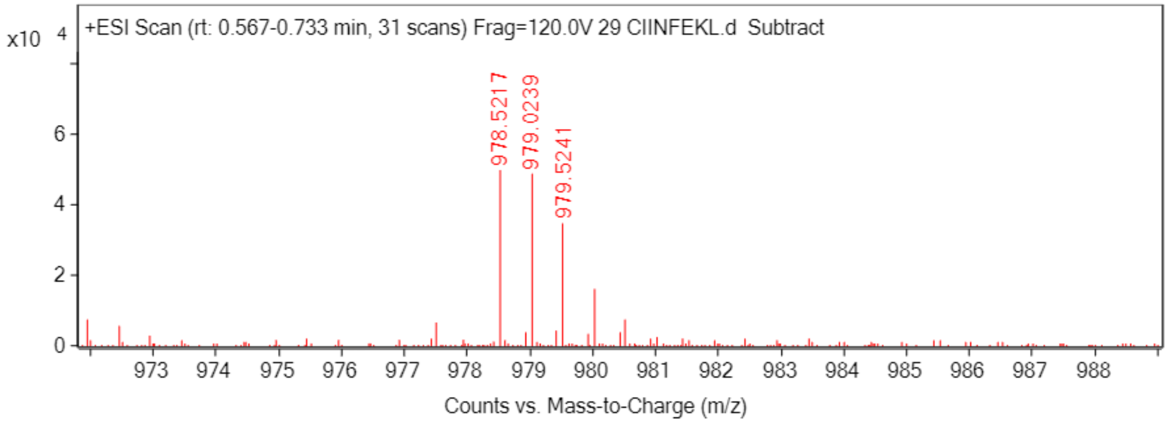

**Scheme S6. Synthesis CSIINFEKL**

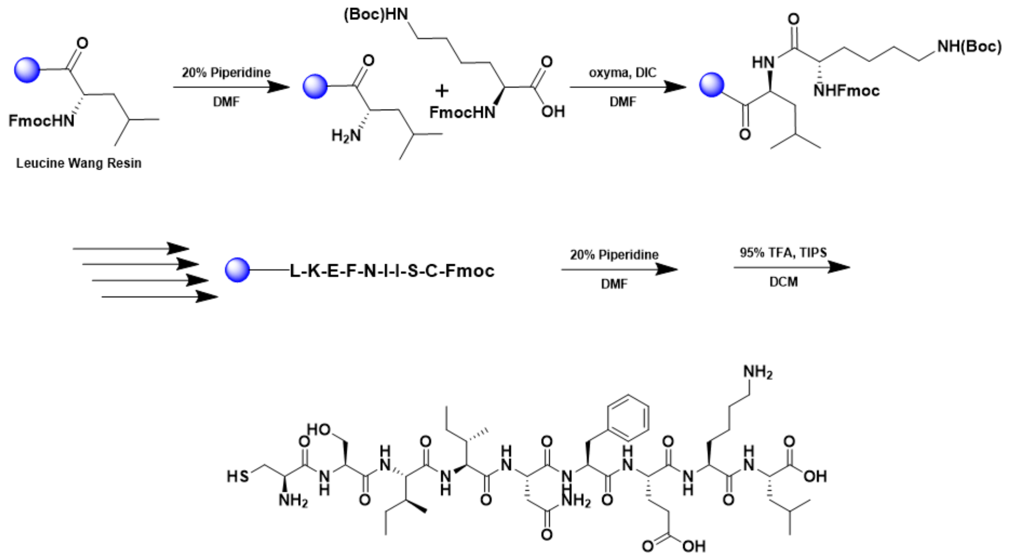


A 25 mL vessel of CEM discover bio manual peptide synthesizer was charged with 0.25 mmol of leucine wang resin. The Fmoc group was removed by using a 20% piperidine solution in DMF (10 mL). Using Synergy software, the deprotection protocol was run. The piperidine solution was drained and the resin was washed with DMF (4 x 10 mL). Fmoc-L-lysine(Boc)-OH (5 eq, 1.25 mM) along with Oxyma (5 eq, 1.25 mM) and DIC (5 eq, 1.35 mmol) in DMF was added to the reaction vessel and the coupling protocol was run. The amino acid solution was drained, and the resin was washed with DMF (2 x 10 mL). The fmoc removal and coupling procedure was repeated as before using the same equivalencies for the remaining amino acids. To remove the peptide from resin, a TFA cocktail solution (95% TFA, 2.5% TIPS, and 2.5% DCM) was added to the resin and agitated for 2 hours. The resin was filtered, and the resulting solution was concentrated in vacuo. The peptide was triturated with cold diethyl ether and purified using reverse phase HPLC using $H_2O/CH_3CN$. The sample was analyzed for purity using a Waters 1525 Binary HPLC Pump using a Phenomenex Luna 5u C8(2) 100A (250 x 4.60 mm) column; gradient eluted with $H_20/CH_3CN$. Molecular weight was confirmed using high resolution electrospray ionization mass spectrometry (HRMS, ESI/MS) analyses obtained on an Agilent 6545B Q-TOF LC/MS equipped with 1260 infinity II LC system with auto sampler. The final peptide product was lyophilized and stored at -20$^{o}$C until further use.

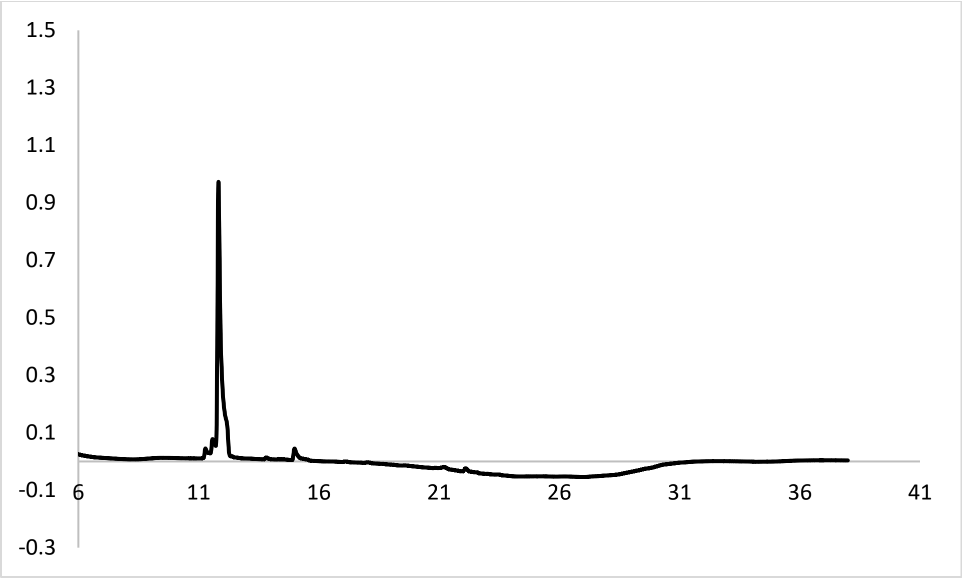


ESI-MS calculated [M+H$^+$]: 1066.5602, found 1066.5612.

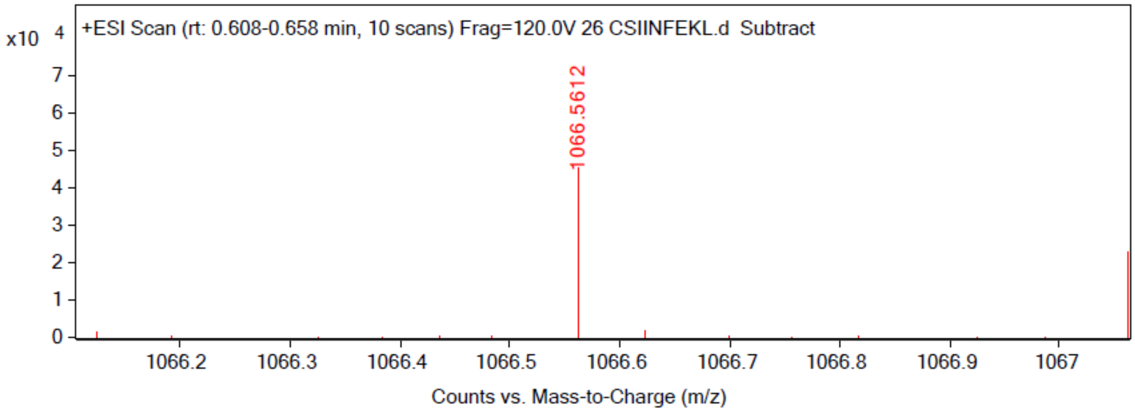

**Scheme S7. Synthesis SIINFKELC**

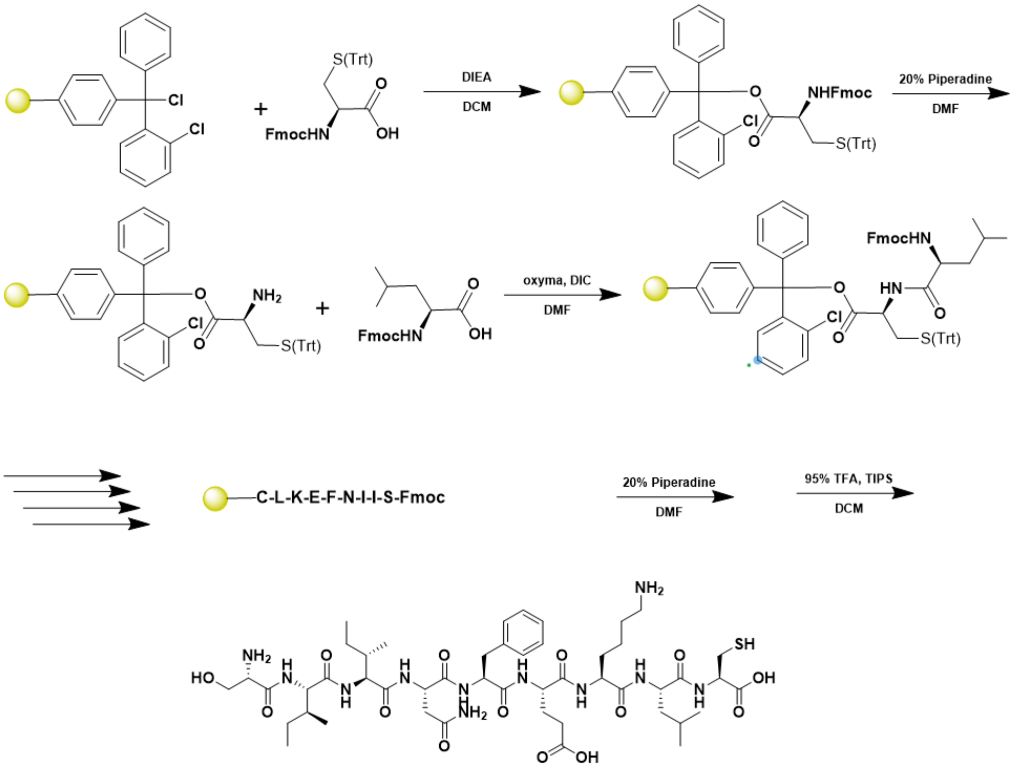


A 25 mL peptide synthesis vessel charged with 2-chlorotrityl chloride resin (0.25mmol) was added Fmoc-L-Cys(Trt)-OH (1.1 eq, 0.275 mmol) and DIEA (3 eq, 0.75 mmol) in dry DCM. The resin was agitated for 1 h at ambient temperature and washed with MeOH and DCM (3 x each). The Fmoc group was removed by using a 20% piperidine solution in DMF for 30 min at ambient temperature, then washed as before. Fmoc-L-leucine-OH (5 eq, 1.25 mM) along with Oxyma (5 eq, 1.25 mM) and DIC (5 eq, 1.35 mmol) in DMF was added to the reaction vessel and agitated for 2 h at ambient temperature. The Fmoc removal and coupling procedure was repeated as before using the same equivalencies for the remaining amino acids. To remove the peptide from resin, a TFA cocktail solution (95% TFA, 2.5% TIPS, and 2.5% DCM) was added to the resin and agitated for 2 hours. The resin was filtered, and the resulting solution was concentrated in vacuo. The peptide was triturated with cold diethyl ether and purified using reverse phase HPLC using $H_2O/CH_3CN$. The sample was analyzed for purity using a Waters 1525 Binary HPLC Pump using a Phenomenex Luna 5u C8(2) 100A (250 x 4.60 mm) column; gradient eluted with $H_20/CH_3CN$. Molecular weight was confirmed using high resolution electrospray ionization mass spectrometry (HRMS, ESI/MS) analyses obtained on an Agilent 6545B Q-TOF LC/MS equipped with 1260 infinity II LC system with auto sampler. The final peptide product was lyophilized and stored at -20$^{o}$C until further use.

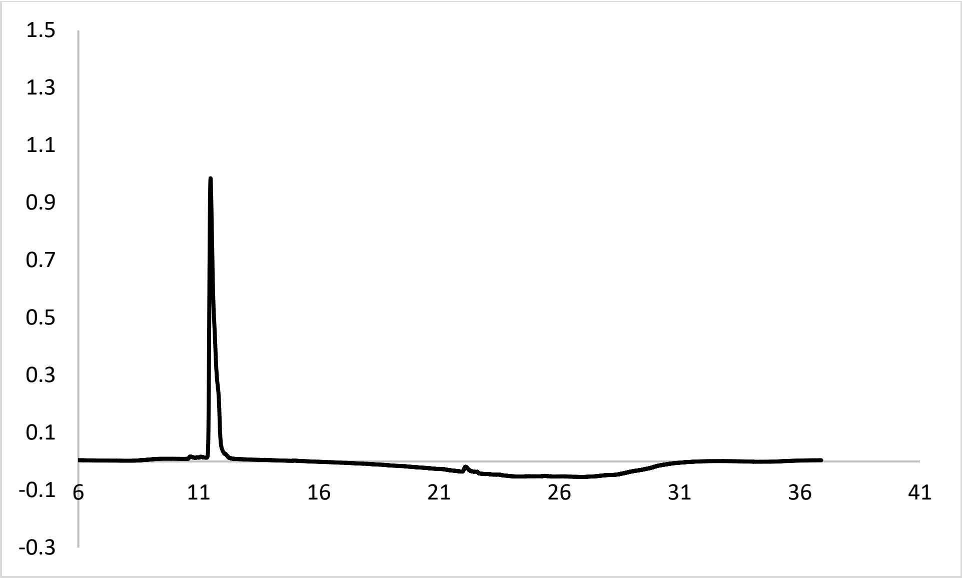


ESI-MS calculated [M+H$^+$]: 1066.5602, found 1066.5560.

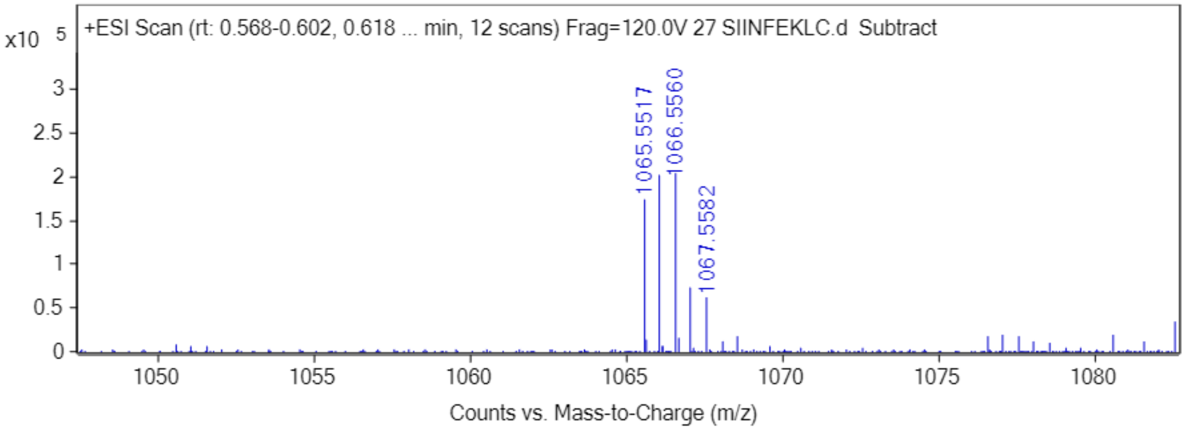

Scheme 8: **Synthesis of SNFVSAGI**

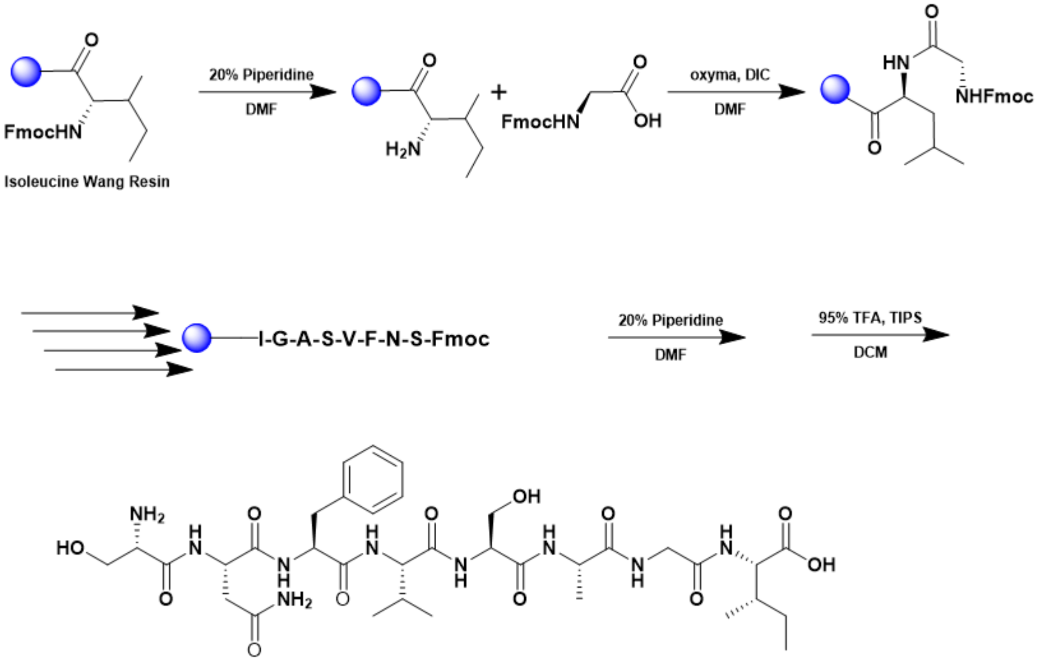


A 25 mL vessel of CEM discover bio manual peptide synthesizer was charged with 0.25 mmol of isoleucine wang resin. The Fmoc group was removed by using a 20% piperidine solution in DMF (10 mL). Using Synergy software, the deprotection protocol was run. The piperidine solution was drained and the resin was washed with DMF (4 x 10 mL). Fmoc-L-glycine (5 eq, 1.25 mM) along with Oxyma (5 eq, 1.25 mM) and DIC (5 eq, 1.35 mmol) in DMF was added to the reaction vessel and the coupling protocol was run. The amino acid solution was drained, and the resin was washed with DMF (2 x 10 mL). The fmoc removal and coupling procedure was repeated as before using the same equivalencies for the remaining amino acids. To remove the peptide from resin, a TFA cocktail solution (95% TFA, 2.5% TIPS, and 2.5% DCM) was added to the resin and agitated for 2 hours. The resin was filtered, and the resulting solution was concentrated in vacuo. The peptide was triturated with cold diethyl ether and purified using reverse phase HPLC using $H_2O/CH_3CN$. The sample was analyzed for purity usinga a Waters 1525 Binary HPLC Pump using a Phenomenex Luna 5u C8(2) 100A (250 x 4.60 mm) column; gradient eluted with $H_20/CH_3CN$. The final peptide product was lyophilized and stored at -20$^oC$ until further use.

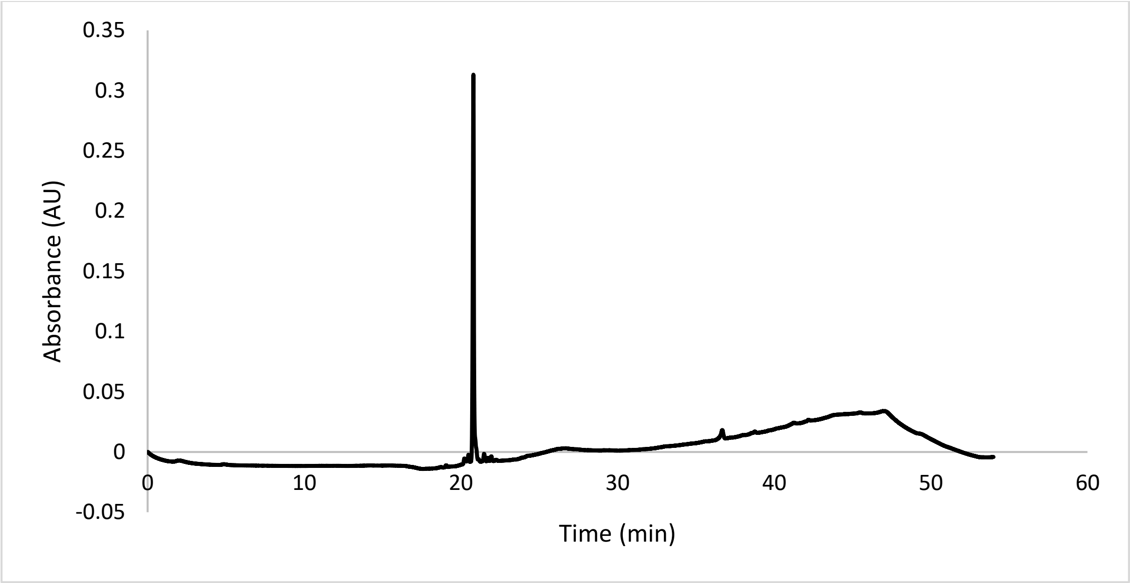


ESI-MS calculated [M+H$^+$]: 794.4043, found 794.4072.

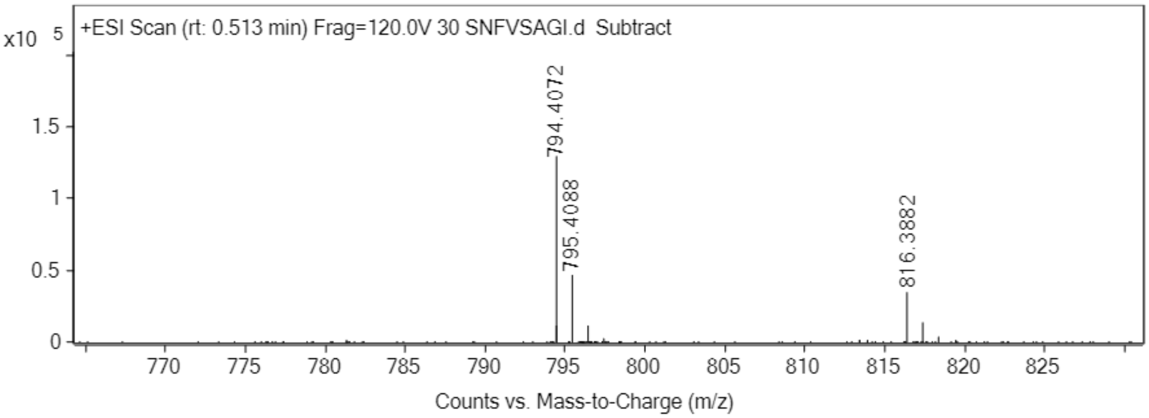
